# KIF5B drives meiotic chromosome dynamics via interaction with the KASH5-LINC complex

**DOI:** 10.1101/2025.05.28.656678

**Authors:** Y. Ditamo, W. Shi, L. Previato, J. P. Gillies, A. Carbajal, K. P. Nowak, M. L. Marin, M. Kinter, M. E. DeSantis, C. G. Bisig, R. J. Pezza

## Abstract

Telomere-led rapid prophase chromosome movements (RPMs) during meiotic prophase are critical for homologous chromosome pairing and proper meiotic progression. These movements are generated by the cytoskeleton and are transmitted to the telomeres via the LINC complex, yet the cytoplasmic components that generate these forces remain poorly defined. Among candidates of microtubule-associated motor proteins in mouse primary spermatocytes, we confirmed KIF5B as a specific interactor of the KASH5-LINC complex. Total internal reflection fluorescence microscopy and microtubule sedimentation assays performed with purified recombinant proteins suggest a direct interaction between KASH5 and KIF5B on microtubules, enhanced by MAP7, a known KIF5B-recruiting and activating cofactor. Mapping the KIF5B-binding surface of KASH5 revealed that KASH5 N-terminal EF-hand domains mediate the interaction. Further, in vivo KIF5B-KASH5 interaction and KIF5B role in RPMs are evidenced as (1) KIF5B is recruited by KASH5-SUN1 to the nuclear envelope in two different cultured somatic cell models, (2) KIF5B is telomere-associated and colocalizes with KASH5, and microtubules associated with the nuclear envelope in mouse spermatocytes, and (3) chemical inhibition of KIF5B reduces telomere-led chromosome motions. Altogether, our findings identify the KIF5B kinesin as a previously unrecognized component of the force-generating machinery that drives chromosome movement during meiotic prophase I, acting through KASH5 as a specific nuclear membrane adaptor.

## Introduction

Homologous chromosome pairing is necessary for the proper segregation of meiotic chromosomes at the first meiotic anaphase, and failure to accomplish these processes results in phenotypes ranging from sterility to aneuploid offspring with developmental abnormalities (*1–5*).

Chromosome interactions and meiotic recombination in prophase are accompanied by telomere-led chromosome movements, also known as rapid prophase movements (RPMs), that are widely conserved in organisms from yeast to mammals (*6–24*). RPMs have been proposed to assist homologous chromosome interactions (e.g., homologous chromosomes pairing and synapsis) (*20, 25–32*), reduce and/or resolve heterologous interactions (*9, 16, 26, 27, 33, 34*), and help resolve chromosome interlocks (*35, 36*). RPMs have been connected to the transitional clustering of chromosome ends, ultimately leading to the “meiotic bouquet” chromosome configuration. At least in some organisms, the meiotic bouquet has been proposed to promote initial homologous connections leading to homologous chromosome pairing (*11, 14, 37–45*). Together, the data point to the importance of RPMs in critical meiotic events required for the normal progression of gametogenesis.

RPMs are driven by the connection of telomeres to the cytoskeleton via protein bridges through the intact nuclear envelope termed the LINC complex (*46*). In the mouse, the LINC complex consists of inner nuclear envelope SUN (<u>S</u>ad1-<u>UN</u>C84) domain proteins and outer nuclear membrane KASH (*<u>K</u>larsicht,* <u>A</u>nc-1, and <u>S</u>yne <u>h</u>omology) domain proteins. Telomeres associate with the intranuclear domain of the SUN proteins, and the extranuclear portion of the KASH protein connects to the cytoskeleton (reviewed in: (*47–49*)).

In mammals, worms, fission yeast, and plants, RPMs require microtubules (*11, 13, 20, 21, 31, 32, 34, 50–52*). Associated with microtubules, dynein is the only molecular motor so far suggested to act as a component of the force-generating RPMs in mammals (*11, 31, 53–55*). In *Arabidopsis thaliana,* normal levels of meiotic RPMs depend on the kinesin motor protein PSS1, which is recruited to the nuclear envelope by KASH-domain protein SINE3 (*56*). These incisive studies have provided the impetus to re-examine the participation of diverse cytoplasmic components of the machinery generating RPMs in mammals.

In this work, we use proteomic approaches to screen and identify kinesins associated with microtubules in mouse spermatocytes active in RPMs. We then used biochemical, cytological, and functional approaches to characterize the KIF5B motor protein associated with the KASH5-LINC complex. Our work identified KIF5B as a novel extranuclear component of the machinery that interacts with the LINC to generate RPMs. Defective meiotic chromosome dynamics invariably lead to the generation of aneuploid gametes; the new insights we provide here on fundamental mechanisms required for proper chromosome dynamics represent a significant advance in understanding how normal meiotic progression is achieved and how its disruption may lead to disease.

## Results

### Identification of motor proteins associated with microtubules in meiotic prophase spermatocytes

Microtubules and microtubule-associated proteins are of critical importance for RPMs activity in mammals, worms, fission yeast, and plants. Yet components of the extranuclear portion of the machinery that promotes RPMs have remained understudied. Here, we undertook a proteomics-based screen to identify motor proteins bound to microtubules in mouse spermatocytes at meiotic stages, when RPMs are active. To do so, we performed microtubule purification and enrichment of associated motor proteins using mouse seminiferous tubules (Fig. 1A) (*57*) followed by protein identification by mass spectrometry (Fig. 1B and Table S1). We used testis from two-month-old mice containing all stages of gamete development from spermatogonia to spermatozoa (*58*). We contrasted these samples with those obtained from 13-day-old mice, in which seminiferous tubules are populated only with primary spermatocytes, with RPMs at their maximum (*11*). Comparison of proteins observed across these samples should allow us to identify microtubule-associated molecular motors enriched in the 13-day-old testis spermatocytes, where RPMs are prominent (*11*). Figure 1B shows kinesins and dynactin enriched in the 13-day-old testis. In the graph, proteins to the right of the dotted line have higher abundance (summed Mascot score) in 13-day-old mice than in adult mouse testes. Specifically, we observed high signal for members of the kinesins superfamily of molecular motors (KIF5B, KIF11, KIF13B, KIF5C, KIF2A, KIF21A, KIF15, KIF5A, KLC2, and KLC4) in all analyzed samples (Fig. 1B, Table S1). In agreement with previous reports, we also identified dynein and dynactin (*11, 31, 54, 55*). We observed DYNC1H1, DCTN1, DYNC1I2, DCTN2, DYNC1LI2, DYNC1LI1, DNAAF1, DCTN4, DYNC1I1, and DYNLRB1, which provide a detailed description of dynein and dynactin subunits that may participate in RPMs, surpassing that previously described (*11, 31, 53*). The results obtained from 13-day-old mouse testis, in which most spermatocytes are at the zygotene stage (Fig. 1B), suggest that the motors we identified associated with microtubules are present at spermatocyte stages when RPMs are prominent. Consistent with the proteomic data, the presence of KIF5B - one of the most abundant motors detected in our screen - was independently confirmed by western blot analysis (Fig. 1C).

**Figure 1.**
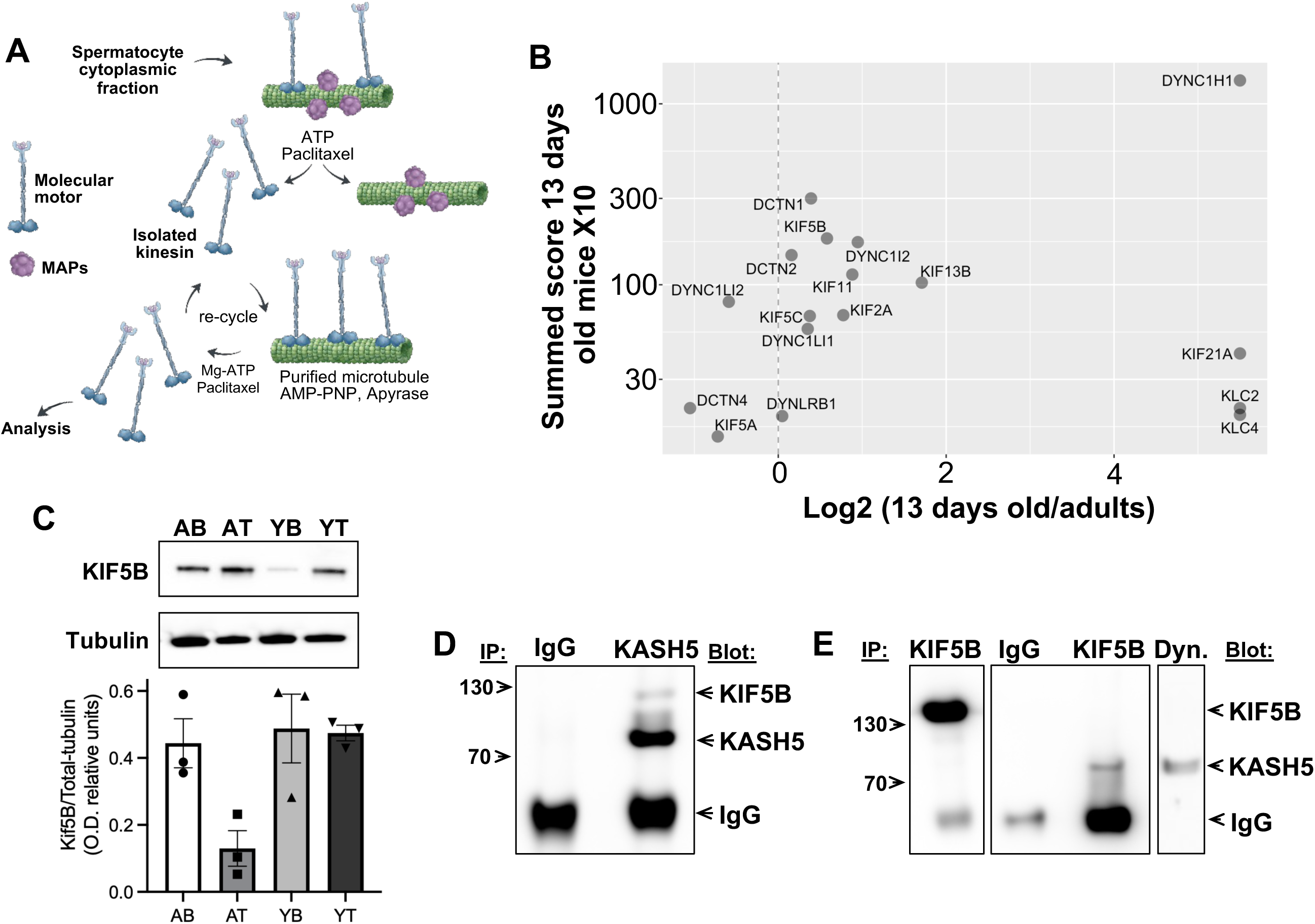
KASH5 interacts with KIF5B in mouse spermatocytes undergoing meiosis. (A) Schematic of the strategy used to enrich microtubule-associated proteins in testis homogenates from adult and 13-day-old wild-type (n = 3, pooled). (B) Graph showing representative kinesin and dynein family members identified by mass spectrometry after purification using the protocol depicted in A. The graph shows a comparative analysis of wild type 13-day-old testis versus adult testis (6-week-old). (C) Microtubule-associated KIF5B purified as shown in (A) from AB, adult brain; AT, adult testes; YB, young brain; and YT, young testes were analyzed by western blots to confirm the presence of KIF5B. Quantitation of KIF5B versus total tubulin immunosignal in 3 independent experiments is also shown; bars are standard error. (D) Immunoprecipitation with a KASH5 antibody or control unspecific IgG was analyzed by sequential immunoblotting with KASH5 and then KIF5B antibodies. Note that KASH5 pulldown itself and KIF5B, whereas IgG does not. For each condition, we used n = 6 testes from 3 mice. The experiment was performed twice. (E) Immunoprecipitation with control IgG, KIF5B, or Dynein (Dyn.) antibody was analyzed by immunoblotting with KIF5B (left panel) and KASH5 antibodies (mid and right panels). Note that KIF5B can pull down itself and KASH5, n = 6 testes from 3 mice, pooled.

Our results are consistent with a model in which KIF5B, together with previously identified dynein (*11*), participates in RPMs of mouse spermatocytes. Nevertheless, we cannot exclude the possibility that some of the molecular motors identified come from a minor population of somatic cells present, or that, in prophase I spermatocytes, they may participate in alternative functions, such as organelle transport.

### KIF5B interacts with the LINC complex via KASH5 in mouse spermatocytes

If KIF5B participates in RPMs, we would expect it to interact with KASH5, a component of the LINC complex located in the outer nuclear membrane and extending into the cytoplasm. To test this hypothesis, we performed immunoprecipitation of KASH5 from total extracts of 13-day-old testes and analyzed the co-immunoprecipitated proteins by western blotting. We observed that, in contrast to nonspecific control rabbit IgG, KASH5-specific antibodies efficiently immunoprecipitated KASH5 and co-immunoprecipitated KIF5B from testis extracts (Fig. 1D).

In complementary experiments, we confirmed that KIF5B-specific antibodies, but not nonspecific IgG, immunoprecipitated KASH5 from total testis extracts (Fig. 1E). Dynein has been associated with RPMs in mouse spermatocytes (*11*). As expected, KASH5 was also detected in Dynein immunoprecipitates (Fig. 1E).

Together, these results demonstrate that KIF5B interacts with KASH5, suggesting that KIF5B forms part of the molecular machinery responsible for generating RPMs in mouse spermatocytes.

In our studies, KIF5B is among the most abundant motor proteins detected in spermatocytes with active RPMs. We also observe a strong and consistent association of KIF5B with both KASH5 and microtubules. Together, these findings identify KIF5B as a strong candidate for a kinesin involved in RPMs. Accordingly, the subsequent sections focus primarily on KIF5B when examining kinesin function.

### KASH5 directly interacts with KIF5B, which is associated with microtubules

To directly test whether KASH5 could form a complex with and function as an adaptor of KIF5B, we monitored the association between the two proteins using a total internal reflection fluorescence (TIRF)-based single-molecule in vitro binding assay. First, we confirmed that the recombinant, purified full-length KIF5B bound tightly to microtubules in the presence of AMP-PNP, a non-hydrolyzable ATP analog (Fig. 2A). KIF5B showed a substantial reduction in binding to microtubules in the presence of ATP. This is expected as an active KIF5B hydrolyzes ATP, resulting in a higher microtubule dissociation constant (Fig. 2A).

**Figure 2.**
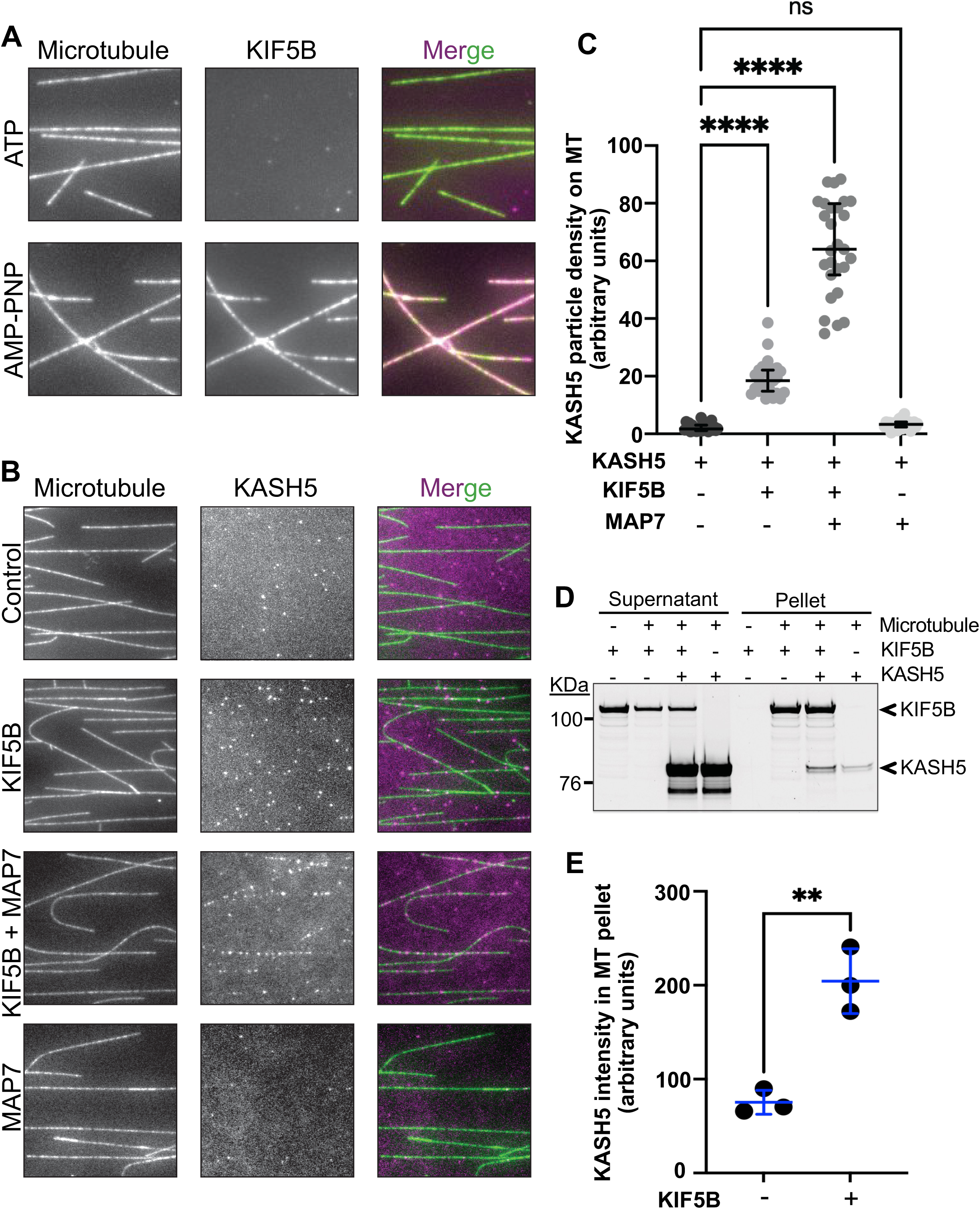
KIF5B increases KASH5 association with microtubules. (A) TMR-labeled KIF5B incubated with HiLyte 488-labeled microtubules in the presence of ATP or AMP-PNP. (B) TMR-labeled KASH5 incubated with HiLyte 488-labeled microtubules in the presence or absence of KIF5B and/or MAP7 under the condition of AMP-PNP. (C) Quantification of KASH5 particle intensity on microtubules (MT) in the absence or presence of unlabeled KIF5B and/or MAP7 under the condition of AMP-PNP, as shown in (B). KASH5 particle intensity is defined as the number of KASH5 particles per mm2 of microtubule area. Each data point represents a single field of view (field size: 81.92 mm2). Statistical analysis was performed using ordinary one-way ANOVA with Dunnett’s multiple comparisons test (**** equals P < 0.0001). Number of replicates, n = 3. (D) SDS-PAGE gel showing supernatant and pellet fractions from the microtubule pelleting assay. (E) Quantification of KASH5 intensity in the pellet fraction from the microtubule pelleting assay shown in (D). Statistical analysis was performed using an unpaired t-test (** equals P < 0.01). Number of replicates, n = 3.

Next, we asked if full-length KIF5B could recruit TMR-labeled KASH5 to the microtubule surface. In the absence of KIF5B, very little KASH5 colocalized with the microtubule. However, in the presence of KIF5B, we observed a significantly increased level of KASH5-microtubule colocalization (Fig. 2B and C). The KIF5B-dependent KASH5 microtubule binding was further enhanced by the addition of the kinesin cofactor MAP7, which has been shown to promote kinesin-microtubule interaction (*59*). Indeed, inclusion of MAP7 resulted in a greater than 2-fold increase in the number of KASH5 molecules that specifically localized to the microtubule track. In the absence of KIF5B, MAP7 was unable to recruit KASH5 to the microtubule surface (Fig. 2B and C). As MAP7 likely destabilizes the autoinhibited conformation of kinesin, these results could suggest that KASH5’s binding site is partially occluded when KIF5B is autoinhibited (*59*).

We also assessed KiF5B’s ability to recruit KASH5 to microtubules using a sedimentation-based experiment. Here, co-sedimentation of KASH5 with polymerized microtubules was assessed in the absence and presence of KIF5B. In confirmation of the TIRF-based binding assay, the presence of KASH5 in pelleted microtubules substantially increased in the presence of KIF5B (Fig. 2D and E).

In sum, TIRF-microscopy-based assays and microtubule co-sedimentation experiments support a model in which KASH5 directly interacts with KIF5B and acts as a newly identified kinesin adaptor.

### KASH5 N-terminal EF-hand domains are essential for the interaction with KIF5B

To further understand the KASH5-KIF5B interaction and potential cooperation in RPM generation, we performed direct yeast-two-hybrid assays (Fig. 3 and Fig. S1). We first confirmed that mouse KASH5 (70-579 amino acids) as a bait interacts with mouse KIF5B (1-963 amino acids) as a prey (Fig. 3A). We then used KASH5 70-579 amino acids and different N-termini and a C-terminus truncated versions of mouse KASH5 as bait to monitor interaction with 1-963 amino acids of KIF5B. Interaction with KIF5B apparently required amino acids 70-156 of KASH5 (Fig. 3B), a region of the N-terminus likely residing in the cytoplasm. Analysis of the KASH5 N-termini amino acid sequence reveals a conserved protein portion predicted to have two consecutive helix-loop-helix EF-hand domains (Fig. 3C). The EF-hand domain is a structure where a calcium ion is coordinated by a loop region connecting two alpha helices (*60*). Although it is apparently not regulated by cellular calcium levels (*55*), KASH5’s EF-hands domains have been shown to be essential in the interaction with dynein, as KASH5-dynein interaction is disrupted by specific mutations in this protein region (*54, 55*). In our experiments, deletion of the 70-156 region, which disrupts interaction with KIF5B, removes both helices and the EF-hand domain 1, as well as the incoming helix associated with the EF-hand domain 2, and approximately 50% of the residues predicted to form EF-hand domain 2. These results implicate the two EF-hand domains of KASH5 in the interaction with KIF5B.

**Figure 3.**
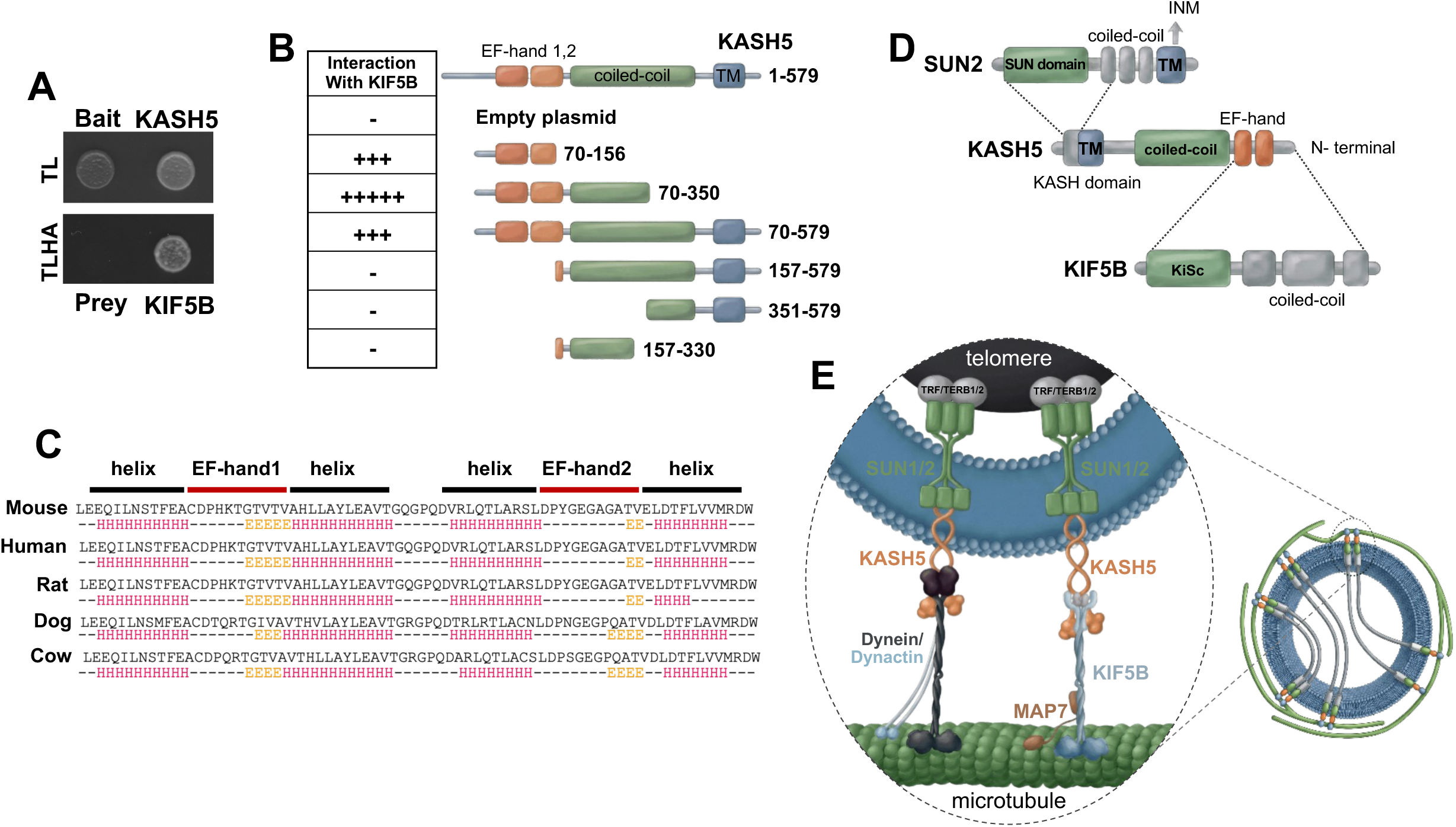
The EF-hand N-terminal domain of KASH5 is implicated in KIF5B interaction. (A) mouse KASH5 interacts with mouse KIF5B in a direct yeast-two-hybrid assay. (B) Study of KASH5 domain interaction by direct yeast-two-hybrid analysis. The schematic represents the predicted functional domains of KASH5, and truncation mutants created to map sites of KIF5B-KASH5 interaction. The negative (−) and positive (+) symbols represent the absence or presence of interaction. Five positive symbols represent maximum interaction strength. See Fig. S1 for details. (C) Protein sequence alignment of KASH5 in different species and predicted secondary structure. The helix and EF-hand domains are indicated. (D) Schematic depicting details of SUN2-KASH5-KIF5B interaction found in this and previous studies. (E) Schematics representing the proposed model for meiotic chromosome telomere-nuclear envelope attachment and connection to kinesin and dynein on microtubules. A schematic representation of a zygotene-pachytene spermatocyte nucleus is also shown.

We conclude that KIF5B N-terminal EF-hand1 and EF-hand2 domains are required for the interaction with KIF5B. Together with previous work showing that protein regions in SUN2 and KASH5 are responsible for their interaction (*61, 62*), our results help construct a model for the KIF5B-LINC-KASH5 complex, proposed to generate the primary forces driving RPMs in mouse spermatocytes (Fig. 3 D and E).

### KASH5 can recruit KIF5B to the nuclear envelope

To assess whether the interaction between KASH5 and KIF5B can recruit KIF5B to the nuclear envelope, we utilized an established immunofluorescence-based molecular motor relocalization assay developed in HeLa cells (*54, 63*). We transfected these cells with vehicle control, SUN1-myc only, or SUN1-myc plus two alternative FLAG-KASH5 constructs (Fig. 4A-D and Fig. S2A and B). Cell cycle analysis revealed that most cells were in S phase in our experiments, and no changes were observed after cell transfection (Fig. S2C). HeLa cells have detectable endogenous levels of KIF5B and SUN1, and, as KASH5 is a meiosis-specific protein, there is no endogenous KASH5. KASH5 localization at the nuclear membrane depends on SUN1, which may be limited by the lower endogenous levels of SUN1. For this reason, ectopic SUN1 overexpression is used to achieve levels comparable to those of the FLAG-KASH5 constructs (*54*).

**Figure 4.**
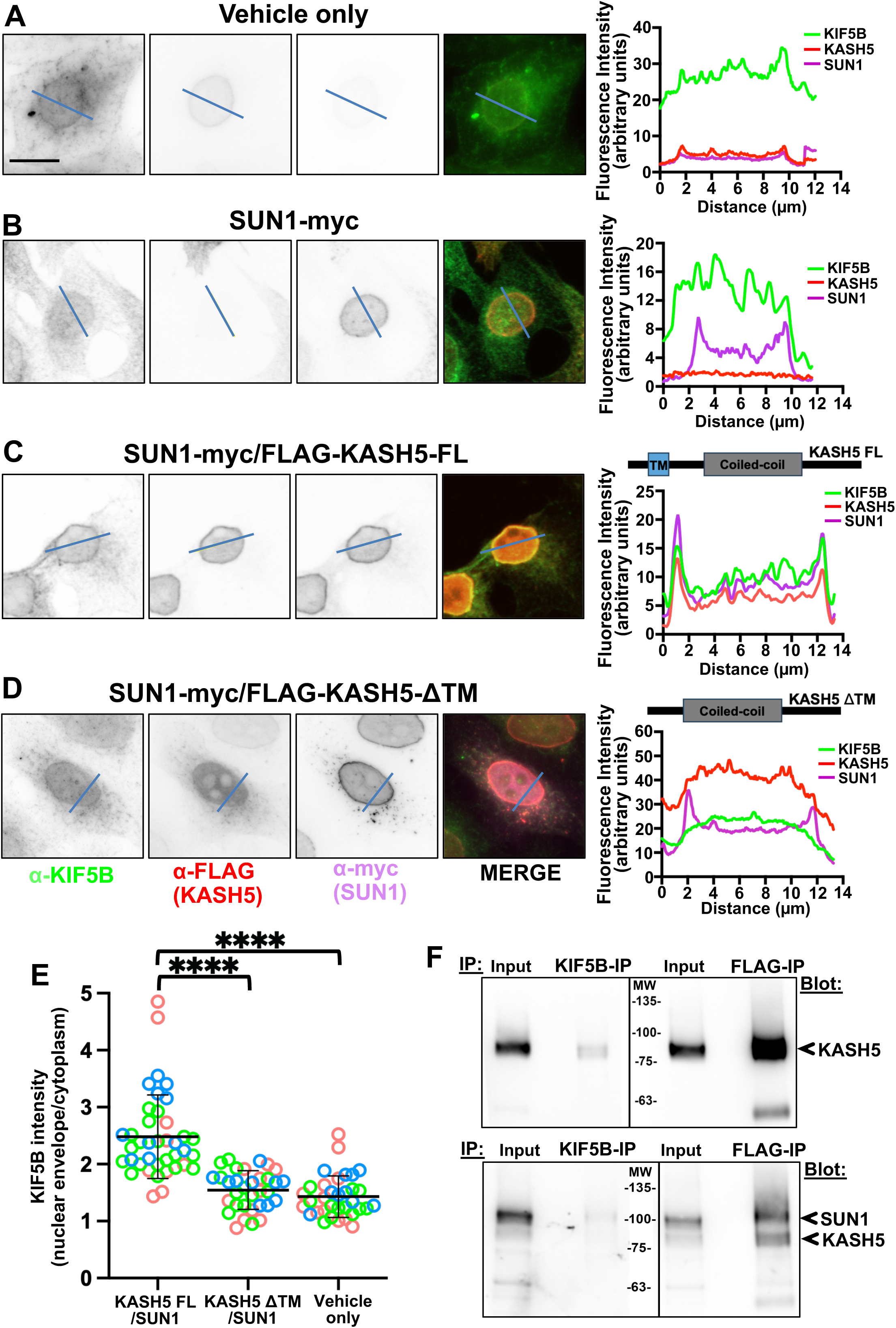
KASH5-mediated recruitment of KIF5B to the nuclear envelope of HeLa cells. Immunofluorescence of HeLa cells transiently transfected with vehicle only (A) or SUN1-myc only (B), SUN1-myc and FLAG-tagged KASH5-FL (full length) (C), or SUN1-myc and FLAG-tagged KASH5-ΔTM (missing the transmembrane domain) (D) constructs and analyzed using anti-KIF5B, anti-FLAG (KASH5), and anti-myc antibodies (SUN1). Scale bar = 10 µm. Light blue lines on the images indicate where a line-scan plot was performed, as shown on the right. (E) Quantitation of KIF5B fluorescence intensity in three experimental conditions. Each data point represents the ratio of NE/cytoplasmic intensity for KIF5B or KASH5 in a single cell from three biological replicates (represented in different colors). Number of cells quantified, n = 39, 33, and 34 for KASH5-FL, KASH5-ΔTM, and non-transfected (vehicle only), respectively. The Kruskal-Wallis test was performed for every pairwise comparison. (F) Western blots showing the results of immunoprecipitation using either a KIF5B antibody or an anti-FLAG (KASH5) antibody, followed by immunoblotting with an anti-FLAG antibody. Note that KIF5B pulls down KASH5, and KASH5 immunoprecipitates itself (upper panel). The lower panel shows proteins precipitated with a KIF5B antibody or an anti-FLAG (KASH5) antibody, followed by immunoblotting with an anti-myc (SUN1) antibody. Note that SUN1 co-immunoprecipitated with KIF5B and KASH5. Inputs (1.5 % of total lysate) and molecular weight markers are also shown.

As expected, the distribution of endogenous KIF5B was relatively diffuse throughout the cytoplasm with no distinct nuclear envelope signal in cells when neither SUN1 nor KASH5 was transiently transfected (vehicle only, Fig. 4A) or when cells were transfected with SUN1-myc only (Fig. 4B).

We observed a well-defined nuclear ring of KIF5B discernible in most cells expressing moderate levels of both SUN1-myc and FLAG-KASH5-FL (full length) (Fig. 4C and Fig. S2A), but not in cells expressing SUN1-myc plus FLAG-KASH5-ΔTM (the latter, which deletes the KASH5 membrane interaction domain) (Fig. 4D and Fig. S2B).

We evaluated protein enrichment at the nuclear envelope by measuring pixel intensity along a scan line drawn across a nuclear cross-section (blue line in Fig. 4A-D and Fig. S2A and B). We then generated fluorescence intensity profiles for each fluorescent channel (Fig. 4A-D and Fig. S2A and B), and quantified relative KIF5B fluorescent intensity per cell (Fig. 4E). Indeed, in cells in which SUN1-myc plus FLAG-KASH5-FL were expressed we observed a clear peak of spatial colocalization for KIF5B, SUN1, and KASH5 at the nuclear envelope (Fig. 4C and Fig. S2A) and increased KIF5B fluorescence intensity per cell (Fig. 4E). This is in sharp contrast to that observed in cells with vehicle alone, SUN1-myc alone, or SUN1-myc plus FLAG-KASH5-ΔTM control (Fig. 4 A, B, D, and E and Fig. S2B).

We observed similar KIF5B relocalization to the nuclear membrane after transfection with FLAG-KASH5-FL, but not with FLAG-KASH5-ΔTM, in Neuro 2 mouse cells (Fig. S2D). This shows that KASH5 can also recruit KIF5B to the nuclear envelope in a mouse-based ectopic system.

Co-immunoprecipitation assays in HeLa and Neuro 2 cells confirmed that KIF5B interacts with ectopically expressed FLAG-KASH5 and SUN1-myc in HeLa cells (Fig. 4F and Fig. S3A and B).

We concluded that, in a cultured cell model, the SUN1-KASH5 complex interacts with and recruits KIF5B to the nuclear envelope.

### KIF5B is associated with telomeres and colocalizes with KASH5 and microtubules at the nuclear membrane in mouse spermatocytes

To test the significance of KIF5B-KASH5 interaction in mouse spermatocytes, we immunolocalized KIF5B, KASH5, and microtubules in structurally preserved squashed spermatocytes using confocal microscopy.

We detected significant accumulation of KIF5B associated with KASH5 at telomeres (the latter revealed by the end of SYCP3 linear signal assigned to a chromosome axis) and microtubules associated with the nuclear membrane (Fig. 5A and B). Not all KASH5 at chromosome ends (again, marked by SYCP3) colocalized with KIF5B immunosignal (mean ± SD, 12.8 ± 2.4 % of KASH5/KIF5B colocalization, n = 20 spermatocytes. 13.6 ± 3.1 % of KIF5B/SYCP3 colocalization, n = 20 spermatocytes) (Fig. 5C). This is similar to that we observed for dynein (*11*) and may be explained by the fact that only a subset of telomeres is actively engaged in RPMs through its interaction with the KIF5B-LINC at any given time.

**Figure 5.**
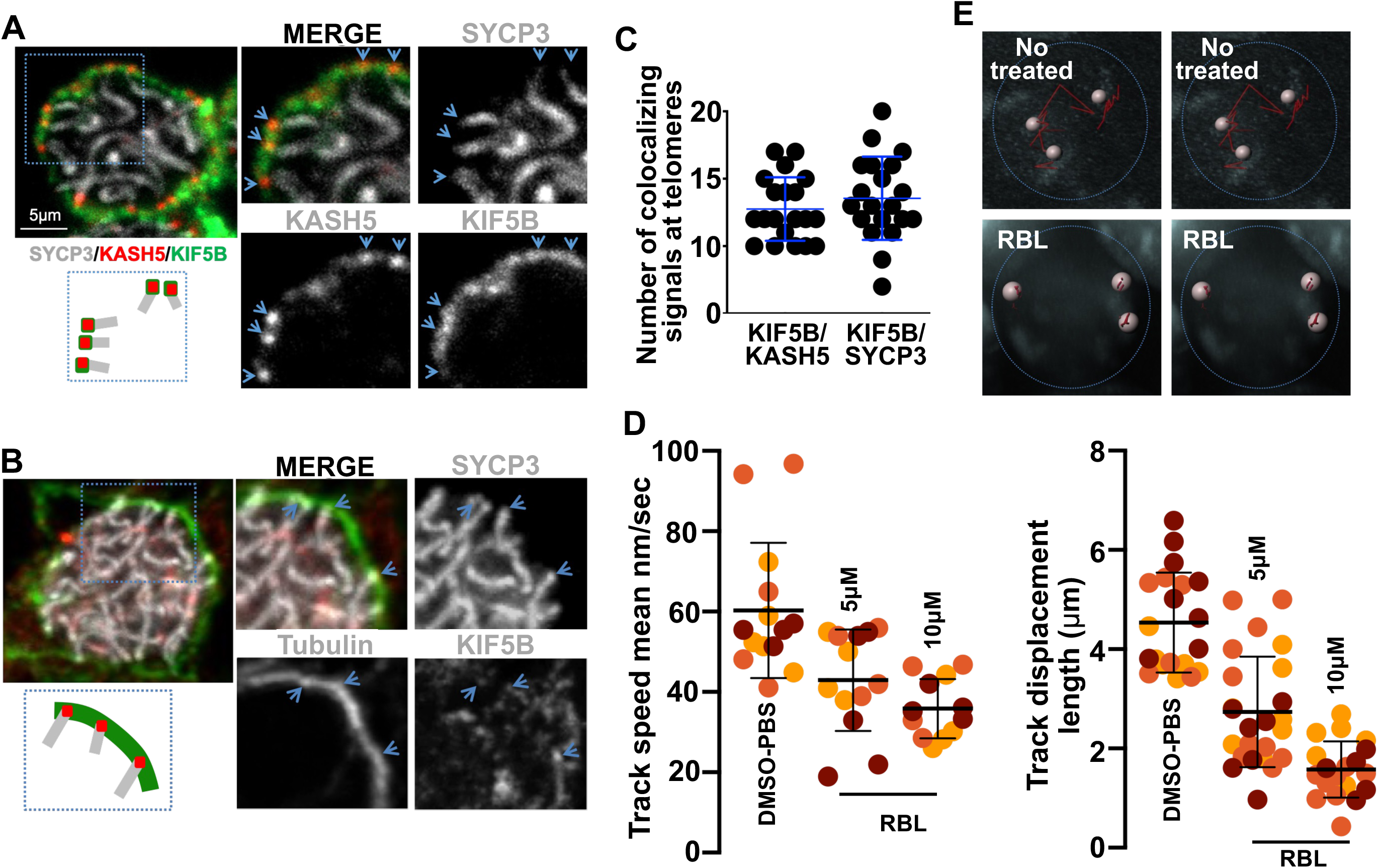
KIF5B colocalizes with KASH5 and tubulin at meiotic chromosomal telomeres and is required for normal levels of RPMs in mouse spermatocytes. Example of wild type pachytene spermatocytes from 4-week-old mice showing localization of components of the synaptonemal complex to mark chromosome cores (SYCP3), KASH5, and KIF5B (A) or chromosome cores (SYCP3), tubulin, and KIF5B (B). The magnified areas highlight the association of the chromosome ends and KIF5B with KASH5 in (A) and KIF5B and tubulin in (B) (arrows). Image representative of three independent immunostaining experiments using spermatocytes obtained from 3 different mice. (C) Quantitation of fluorescent signals for colocalizing KASH5, KIF5B, and SYCP3 shown in A (KIF5B/KASH5 colocalization n = 20 spermatocytes; KIF5B/SYCP3 colocalization n = 20 spermatocytes). Analyzed spermatocytes were obtained from 3 different mice. (D) Quantitation of RPMs track speed mean, and track displacement length in zygotene spermatocytes after seminiferous tubules treatment with 1 % DMSO-PBS (control) or the rose bengal lactone inhibitor (concentrations as indicated); horizontal and vertical lines indicate the median and standard deviation values, respectively. Data were obtained from at least 3 different 6-week-old mice. (E) Representative traces of total spot movements in individual cells, non-treated and treated with role bengal lactone KIF5B inhibitor. Movement in the depicted zygotene spermatocytes was measured for 10 min. Each of the three different representative heterochromatin spots is represented.

Our results show that KIF5B associates with microtubules and with the nuclear envelope-associated protein KASH5 in mouse spermatocytes, suggesting a role for KIF5B in RPMs.

### Inhibition of KIF5B reduces RPMs in mouse spermatocytes

To test the functional requirement of KIF5B in mouse spermatocyte RPMs, we measured the movements of pericentric heterochromatin spots (which mark one end of the acrocentric mouse chromosome complement) in zygotene cells and collected time-lapse 3D image stacks as previously described (*11*).

Compared with control seminiferous tubules incubated with 1% DMSO-PBS (68.75 ± 17.46 nm/second, n = 16 cells obtained from 3 mice), seminiferous tubule explants treated with different doses of the KIF5B inhibitor rose bengal lactone (*64, 65*), which inhibits the interaction between KIF5B and microtubules (*66*), showed a significant decrease in RPM track speed mean (5 µM, 42.92 ± 12.62 nm/s, two tailed t test P < 0.0001, n = 13 cells; 10 µM, 35.83 ± 7.35 nm/s, P < 0.0001, n = 12 cells) (Fig. 5D). Track displacement length, which is the distance between the first and last spot’s position, a measurement of processivity, also decreases significantly when spermatocytes are exposed to the inhibitor (1% DMSO-PBS, 4.53 ± 1.00 μm, two tailed t test P < 0.0001 two tailed t test, n = 20 cells; 5 µM, 2.74 ± 1.11 μm, two tailed t test P < 0.0001, n = 24 cells; 10 µM, 1.58 ± 0.56 μm, two tailed t test P < 0.0001, n = 19 cells) (Fig. 5D).

Figure 5E shows examples of chromosome movement traces in spermatocyte nuclei treated with DMSO-PBS (control) or the indicated doses of KIF5B inhibitor.

The pattern of the microtubule network and the number of KASH5 foci per nucleus remained grossly unaltered after inhibitor treatment (Fig. S4), suggesting that KIF5B *per se*, and not a secondary effect on microtubule network integrity or an effect on the integrity of the LINC complex, accounts for the effects of the inhibitor on RPMs.

Taken together, our results suggest that KIF5B may contribute to generating the mechanical forces that drive RPMs in mouse spermatocytes.

## Discussion

### KIF5B is a structural part of the mechanism generating RPMs in mouse spermatocytes

In this work, we show that the KIF5B motor protein associates with microtubules in spermatocytes at meiotic stages where RPMs are prominent. Furthermore, we demonstrate that KIF5B interacts with the LINC complex component KASH5 and localizes to telomeres associated with microtubules in mouse spermatocytes. Together with our finding that treatment of seminiferous tubules with a KIF5B inhibitor significantly reduces RPM speeds, these results suggest that KIF5B is an integral component of the machinery generating RPMs in mouse spermatocytes.

KIF5B (this work) and dynein-dynactin (*11*) associate with microtubules in prophase I spermatocytes, and inhibition of either dynein (*11*) or kinesins (this work) substantially reduces RPMs, supporting a model in which both dynein and kinesins contribute to the characteristic, complex chromosome movements observed in mouse spermatocytes. It is tempting to propose that the complex movements observed in mouse spermatocytes arise from the dynamic interplay between kinesin- and dynein motors.

Examples of redundant functions for microtubule-sliding motors have been described in spindle elongation during mitotic human cell division (*67, 68*) and in axonal transport in mammalian neurons (*69*). However, this is not always the case. In *Caenorhabditis elegans* hyp7 cells, the KASH(*70*)-domain protein UNC-83 coordinates with both kinesin-1 and dynein at the nuclear envelope to mediate nuclear migration. Although both motor proteins interact with UNC-83, they bind to distinct regions of the protein, suggesting specialized roles. In this case, Kinesin-1 predominantly provides the mechanical force required for nuclear translocation, whereas dynein may serve a regulatory function (*70*). This could also be the case in *Arabidopsis thaliana,* in which the KASH-domain protein SINE3, which recruits the meiosis-specific kinesin PSS1 to the nuclear envelope (*56*).

Our work showing that KIF5B can use meiotic chromosomes as a cargo, with specificity dictated by KASH5 as an adaptor, adds to the number of important cellular activities assigned to different kinesins. For example, KIF11 has been directly implicated in spindle assembly and organization during both meiosis and mitosis (*71, 72*), and KIF13B is primarily involved in intracellular trafficking and cell polarity, with no reported role in spindle-associated regulation in mammalian germ cells (*73*). KIF2A, although associated with the spindle, functions as a microtubule-depolymerizing kinesin (*74*), and KIF21 has been described as a negative regulator of microtubule elongation (*75*).

### KASH5 is a novel kinesin adaptor

Kinesin adaptors are specialized proteins that facilitate the selective recruitment and attachment of cargoes to kinesin motor complexes. By simultaneously binding to specific cargo molecules and to kinesin motor domains or associated light chains, these adaptors ensure the spatial and temporal regulation of intracellular transport. They play a critical role in determining cargo specificity, regulating motor activity, and integrating signaling pathways that modulate motor-cargo interactions in response to cellular cues.

Although adaptor proteins vary widely in sequence, they often share structural features such as an extended coiled-coil domain and distinct regions specialized for cargo interaction (*11, 76–79*). Indeed, we describe that KASH5 binds to KIF5B and exhibits typical structural characteristics of an adaptor, but with a notable unique characteristic: it is a transmembrane protein that connects to and regulates chromosome movements.

Our data provide in vitro and in vivo evidence that KASH5 and KIF5B form a complex and that this complex likely cooperates to promote dynamic chromosome movements in meiotic cells. Specifically, the role of KASH5 as a kinesin adaptor is supported by our findings showing KASH5 colocalization with KIF5B on microtubules, KASH5’s ability to recruit KIF5B to the nuclear envelope, and KASH5-KIF5B direct interaction observed using purified proteins.

KASH5 interacts with both dynein and KIF5B, and these interactions seem to be mediated by the EF-hand domains located in the cytoplasmic N-terminal portion of KASH5. These observations raise interesting questions regarding potential competition and/or redundancy between dynein and kinesins in generating the motor forces that drive complex RPMs in mammals. Supporting this model, several kinesin adaptors have been proposed to function as bidirectional adaptors as they can directly interact with both kinesin and dynein. Notably, their kinesin- and dynein-binding regions often overlap, suggesting that these adaptors might serve as bidirectional switches to coordinate bidirectional transport. For instance, HAP1 and TRAKs have been shown to interact with both dynein and kinesin through the HAP coiled-coil domain (*80–82*). Additionally, some motor adaptors possess regulatory signals, such as binding partners or post-translational modifications, that may enable a controlled switch between dynein- and kinesin-based motility. For example, TRAK proteins interact with Miro, a Ca²⁺ sensor that, upon Ca²⁺ binding, alters the association between TRAK and kinesin, thereby reducing mitochondrial motility (*83, 84*). Similarly, JIP1 contains a phosphorylation site that regulates the transition from kinesin- to dynein-driven transport of APP-positive vesicles (*85*). These bidirectional adaptors help enhance the spatial and temporal control of cargo transport by regulating which motor proteins are active in response to cellular signals and demands.

Collectively, our findings identify the KIF5B-KASH5 complex as an essential component of the molecular machinery driving dynamic chromosome movements in mouse spermatocytes. The localization of KIF5B to both microtubules and telomeres, its direct interaction with KASH5, and the significant reduction in RPMs upon kinesin inhibition, all support a model in which kinesins cooperate with dynein to mediate complex, bidirectional chromosome dynamics during meiosis. In this framework, KIF5B may act as a primary driver of RPMs, providing sustained plus-end-directed motility along microtubules, while dynein generates opposing forces that enable directional switching, fine-tuning, and possibly spatial confinement of chromosome trajectories. The ability of KASH5 to interact with both kinesin and dynein through its EF-hand domains suggests that it may function as a bidirectional adaptor that coordinates or modulates motor activity in response to regulatory signals. We also speculate that this coordinated motor activity is likely influenced by the specific landscape of microtubule post-translational modifications present in spermatocytes. For example, tubulin modifications such as acetylation, detyrosination, or polyglutamylation are known to differentially regulate motor binding, processivity, and force generation, raising the possibility that specific patterns of tubulin post-translational modifications bias motor engagement and tune the balance between kinesin- and dynein-driven motility. Disruption of these regulatory layers could impair chromosome dynamics, and, given the central role of RPMs in homolog pairing, recombination, and faithful chromosome segregation, defects in KIF5B function, its regulation, or the microtubule code may ultimately contribute to meiotic errors, increasing the risk of aneuploidies and associated reproductive disorders in mammals.

## Materials and Methods

### Mice

Mice used in this study were wild-type (C57BL/6)(*86*). Experiments conformed to relevant regulatory standards and were approved by the OMRF IACUC Institutional Animal Care and Use Committee.

### Enrichment of molecular motors from mouse seminiferous tubules

Molecular motors were enriched in 13- and 60-day-old mouse testis, according to (*57*). Three wild type animals were used. Briefly, testes were homogenized in a ratio of 1 g of tissue/1.5 mL of PME buffer [0.1 M PIPES (pH 6.9), 2 mM EGTA, 1 mM MgSO4, and 0.1 mM GTP], and centrifuged at 100,000 X g for 1 h at 4 °C. Supernatant was incubated with paclitaxel 20 μM for 30 min at 37 °C, and then centrifuged at 39,000 g for 30 min at 24 °C to precipitate microtubules. Supernatant was incubated with 2 units/mL apyrase for 30 min at 25 °C. After incubation, an aliquot of freshly prepared MAP-free, paclitaxel-stabilized microtubules (purified according to (*57*)) was added to a final microtubule concentration of 0.1 mg/mL in the presence of 0.5 mM AMP-PNP, and the mixture was incubated for 30 min at 37 °C. Microtubules and bound kinesin were centrifuged at 39,000 X g for 30 min at 4 °C. The pellet was resuspended in 200 μL of PME buffer containing 20 μM paclitaxel and 10 mM Mg-ATP and incubated for 30 min at 37 °C. The released molecular motors were separated from the microtubules by centrifugation at 39,000 × g for 30 min at 4 °C. This preparation was used for mass spectrometry and western blot immunodetection.

### Immunoprecipitation using total mouse testis extracts

Mice testes were homogenized in IP buffer (50 mM Tris-HCl pH 7.5; 150 mM NaCl; 0,1 % IGEPAL; 0,5 % Deoxycholate; 3 mM MgCl2; 3 mM EDTA; 10% Glycerol; supplemented with protease and phosphatase inhibitors). After the homogenization, the cell extracts were incubated on ice for 30 min. The soluble fraction used for immunoprecipitation was then obtained by centrifugation at 50,000 X g for 20 min at 4 °C. For immunoprecipitation, 2 μg of anti-KASH5, KIF5B, KIF2B, or normal IgG antibodies were added to the cytoplasmic extracts, incubated overnight at 4 °C in rotation, followed by incubation with 50 μL of Dynabeads™ M-280 Sheep Anti-Rabbit IgG (Invitrogen #11201D) for 1 h at 4 °C with rotation. Beads were washed four times with ice-cold IP buffer, and bound proteins were eluted by boiling for 5 min with SDS-PAGE sample buffer.

Proteins were separated by 4-15% SDS-PAGE under reducing conditions and either analyzed by mass spectrometry or transferred to nitrocellulose membranes. The blots were probed with individual primary antibodies as indicated and then incubated with HRP-conjugated donkey anti-rabbit antibodies. In all blots, proteins were visualized by enhanced chemiluminescence.

### Mass spectrometry. Protein identification experiment

A GeLC approach was used to identify the proteins in immunoprecipitates. The product of each immunoprecipitation was run approximately 5 cm into an SDS-PAGE gel, and the gel was stained with Coomassie blue. Each lane of the gel was cut into 7 molecular weight fractions. The proteins in each fraction were digested in gel. Briefly, the gel pieces were washed, reduced with DTT (10 mM in 100 mM ammonium bicarbonate), alkylated with iodoacetamide (30 mM in 100 mM ammonium bicarbonate), and digested overnight with trypsin (1 µg per sample in 50 mM ammonium bicarbonate). The peptides produced by the digestion were extracted from the gel pieces, evaporated, and reconstituted for analysis by LC-tandem mass spectrometry to identify the proteins contained in that fraction.

The tandem mass spectrometry system consisted of a Thermo Scientific QEx Plus coupled to an Ultimate 3000 nanoflow HPLC system. The instrument was operated in data-dependent mode, acquiring one full-scan spectrum (resolution = 70,000) and 10 collision-induced dissociation (CID) spectra (resolution = 17,500) per cycle. The LC conditions were a linear 60 min gradient elution from 2 % to 45 % CH_3_CN in 0.1 % formic acid at a flow rate of 150 nL/min. Approximately 20,000 spectra are acquired in each analysis.

Data were analyzed using the program Mascot. The mouse RefSeq database was searched. Basic search parameters included: 1 missed cleavage, 10 ppm precursor mass tolerance, 0.05 Da fragment mass tolerance, dynamic modification oxidized methionine, static modification carbamidomethyl cysteine. Results were exported as an Excel spreadsheet (“original data”, tables S1 and S2) and processed to include all identifications based on 2 or greater peptides. False discovery rates were in the range of 1 %.

Further processing of mascot results was done as follows (“processed_data”, tables S1 and S2): mascot scores found across all fractions within a lane/sample were summed to get the “summed_mascot_score” value. “max_mascot_score” represents the highest mascot score found on a fraction for a given protein within a lane. “Number of entries” represents the number of fractions within a lane in which a given protein was found. For this, we used R (version 4.0.3) (R Core Team, 2020) and the tidyverse package (version 1.3.1, https://doi.org/10.21105/joss.01686). Protein gene symbols were acquired using biomaRt (mapping identifiers for the integration of genomic datasets with the R/Bioconductor package biomaRt(*87*). Plots were done using ggplot2.

### Direct KASH5-KIF5B yeast-two-hybrid analysis

Full-length and truncation fragments of human KIF5B or human KASH5 were cloned into pGADT7-AD (Prey, Clontech), to produce fusions to the Gal4 DNA-binding and activation domains. Plasmids containing full-length KIF5B or KASH5 were constructed by cloning the appropriate PCR products into pGBKT7 (bait, Clontech). All fusions were confirmed by sequencing. Direct two-hybrid assays were conducted in the AH109 strain background. After mating, colonies containing both plasmids were selected using media lacking tryptophan and leucine. Interactions between partners were assayed by growth on synthetic media lacking tryptophan, leucine, adenine, and histidine (see Fig. S1 for references of the interaction intensity). Transformations were carried out according to the manual for the matchmaker kit (BD Biosciences).

### Protein expression and purification

The His-ZZ-TEV-Halo-KASH51-400 was expressed from the pET28 vector backbone in BL21 (DE3) E. coli cells after induction with 0.5LmM Isopropyl ß-D-1-thiogalactopyranoside (IPTG) for 18 hours at 16 °C and 200 rpm. The induced cells were harvested by centrifugation (6,000 X rpm, 20 min, 4 °C). The pellets were resuspended in lysis buffer (30 mM HEPES [pH 7.4], 50 mM potassium acetate, 2 mM magnesium acetate, 1 mM EGTA, 1 mM DTT, 0.5 mM Pefabloc, 10 % [v/v] glycerol) supplemented with 1 X complete EDTA-free protease inhibitor cocktail tablets (Roche). The resuspended cells were incubated on ice for 30 min with 1 mg/mL egg lysozyme and sonicated (50% amplitude, pulse on 5 s, pulse off 25 s). The lysate was clarified by centrifugation (30,000 X rpm, 30 min, 4 °C) in a Type 70Ti rotor (Beckman). The supernatant was collected and incubated with 2 mL of IgG Sepharose 6 Fast Flow beads (Cytiva) equilibrated in lysis buffer for 3 hours with rotation at 4 °C. The beads were collected by centrifugation (1000 × g, 2 min, 4 °C) and resuspended in 2 mL lysis buffer before being transferred to a glass gravity column. Beads were then washed with 100 mL wash buffer (30 mM HEPES [pH 7.4], 200 mM potassium acetate, 2 mM magnesium acetate, 1 mM EGTA, 1 mM DTT, 0.5 mM Pefabloc, 10% [v/v] glycerol) and 50 mL tobacco etch virus (TEV) buffer (50 mM Tris–HCl [pH 8.0], 150 mM potassium acetate, 2 mM magnesium acetate, 1 mM EGTA, 1 mM DTT, 0.5 mM Pefabloc and 10% [v/v] glycerol). The ZZ-TEV tag was cleaved by incubating the beads with TEV protease at a final concentration of 0.2 mg/mL overnight. Cleaved Halo-KASH51-400was concentrated to 1 mL with a 30K molecular weight cut off (MWCO) concentrator (EMD Millipore) and diluted with 1 mL buffer A (30 mM HEPES [pH 7.4], 50 mM potassium acetate, 2 mM magnesium acetate, 1 mM EGTA, 1 mM DTT, 10% [v/v] glycerol). The 2 mL protein sample was loaded into a HiTrapQTM Q HP column (Cytiva) at 1 mL/min. The column was prewashed with 10 CVs of buffer A, 10 CVs of buffer B (30 mM HEPES [pH 7.4], 200 mM potassium acetate, 2 mM magnesium acetate, 1 mM EGTA, 1 mM DTT, 10% [v/v] glycerol), and again with 10 CVs of buffer A. To elute, a linear gradient was run over 26 CVs from 0 to 100% buffer B. The peak fractions were collected and concentrated to 500 μL with a 30K MWCO concentrator (EMD Millipore). The 500 ml protein sample was further subjected to size exclusion chromatography (SEC) on a Superose 6 Increase 10/300 GL column (Cytiva) with GF150 buffer (25 mM HEPES [pH 7.4], 150 mM KCl, 1 mM MgCl2, 1 mM DTT) as the mobile phase at 0.75 ml/min. Peak fractions were collected, supplemented with glycerol to a final concentration of 10 %, concentrated to 0.2–1 mg/mL with a 30 K MWCO concentrator (EMD Millipore), frozen in liquid nitrogen, and stored at –80°C.

The KIF5B-Halo-TEV-ZZ and ZZ-TEV-SNAP-MAP7 plasmids are gifts from Dr. Michael Cianfrocco. PFastBac plasmid containing KIF5B-Halo-TEV-ZZ or ZZ-TEV-SNAP-MAP7 was transformed into DH10EmBacY chemically competent cells with heat shock at 42°C for 15 s, followed by incubation at 37 °C and agitation at 220 rpm for 4 h in S.O.C. media (Thermo Fisher Scientific). The cells were plated on LB-agar plates containing kanamycin (50 μg/ml), gentamicin (7 μg/ml), tetracycline (10 μg/ml), BluoGal (100 μg/ml), and IPTG (40 μg/ml). Cells that contained the plasmid of interest were identified with a blue/white selection after 48–72 hr. Colonies were grown overnight in LB medium containing kanamycin (50 μg/ml), gentamicin (7 μg/ml), and tetracycline (10 μg/ml) at 37 °C and agitation at 220 rpm. Bacmid DNA was extracted from overnight cultures using isopropanol precipitation. 2 mL of Sf9 cells in a well of a 6-well dish at a density of 0.5 × 10^6 cells/mL were transfected with about 2 µg of bacmid DNA using 13 µL of FuGene HD transfection reagent (Promega) and incubated at 27 °C insect cell incubator for 24 hr. The cells were then supplemented with 1 mL of media and incubated at 27°C for 48 hr. Next, the supernatant containing the virus (V0) was harvested by centrifugation at 1000 rpm for 5 min. 1 mL of the V0 virus was used to transfect 50 mL of Sf9 cells at 1 × 10^6 cells/mL to generate the next passage of the virus (V1). Cells were incubated at 27°C for 3 days, and then the supernatant containing the V1 virus was collected for protein expression. To express protein, 4 mL of V1 virus was used to transfect 400 mL of Sf9 cells at a density of 1 × 10^6 cells/mL. Cells were incubated at 27°C for 3 days and harvested by centrifugation (3500 X rpm, 20 min, 4 °C). The pellet was washed with 10 mL of ice-cold PBS and collected again via centrifugation before being flash-frozen in liquid nitrogen and stored at –80°C until needed for protein purification.

To purify KIF5B protein, sf9 cell pellets were thawed on ice and resuspended in 40 mL of kinesin-lysis buffer (30 mM HEPES [pH 7.4], 50 mM potassium acetate, 2 mM magnesium sulfate, 1 mM EGTA, 0.2 mM Mg-ATP, 0.5 mM DTT, 0.1 % octylglucoside, 5 % [v/v] glycerol) supplemented with one complete EDTA-free protease inhibitor cocktail tablet (Roche) per 50 ml. Cells were lysed with a Dounce homogenizer (ten strokes with a loose plunger followed by 15 strokes with a tight plunger). The lysate was clarified by centrifugation (40,000 X rpm, 35 min, 4 °C) in a Type 70Ti rotor (Beckman). The supernatant was collected and incubated with 2 mL of IgG Sepharose 6 Fast Flow beads (Cytiva) equilibrated in kinesin-lysis buffer for 3 hours with rotation at 4 °C. The beads were collected by centrifugation (1000 × g, 2 min, 4 °C) and resuspended in 2 mL lysis buffer before being transferred to a glass gravity column. Beads were then washed with 50 mL low salt wash buffer (30 mM HEPES [pH 7.4], 150 mM potassium acetate, 2 mM magnesium sulfate, 1 mM EGTA, 0.5 mM DTT, 0.1 % octylglucoside, 5 % [v/v] glycerol), 200 mL high salt wash buffer (30 mM HEPES [pH 7.4], 200 mM potassium acetate, 2 mM magnesium sulfate, 1 mM EGTA, 0.2 mM Mg-ATP, 0.5 mM DTT, 0.1 % octylglucoside, 5 % [v/v] glycerol) and 200 mL low salt wash buffer. The ZZ-TEV tag was cleaved by incubating the beads with TEV protease at a final concentration of 0.2 mg/mL overnight. Cleaved proteins in the supernatant were concentrated with a 100 K MWCO concentrator (EMD Millipore) to 500 µl and purified via SEC on a Superose 6 Increase 10/300 GL column (Cytiva) with kinesin buffer as the mobile phase at 0.75 ml/min. Peak fractions were collected, concentrated to 1 mg/mL with a 100 K MWCO concentrator (EMD Millipore), frozen in liquid nitrogen, and stored at −80°C. The purification of MAP7 was similar, except that ATP was not needed for the buffers.

For fluorescent labeling of purified KASH5 or KIF5B, the proteins were mixed with a 10-fold excess of Halo-TMR (Promega) for 10 min at room temperature. Unconjugated dye was removed by passing the protein through a Micro Bio-spin P-6 column (Bio-rad) equilibrated in GF150 buffer supplemented with 10 % glycerol. Small volume aliquots of the labeled protein were flash-frozen in liquid nitrogen and stored at −80°C.

### Single-molecule TIRF microscopy

Microtubule binding assays were performed in flow chambers assembled as described previously38. No. 1-1/2 coverslips (Corning) were functionalized with biotin-PEG by sonication at 40 °C with 100 % EtOH for 10 min, 200 mM KOH for 20 min, and again with 100 % EtOH for 10 min. Coverslips were rinsed with water 3 times in between sonication steps. After the last EtOH wash, the coverslips were incubated overnight in a vacuum desiccation chamber in the dark with methanol containing 5% acetic acid and 1% (3-Aminopropyl) triethoxysilane (Millipore-Sigma). The next day, the coverslips were washed 5 times with water and once with 100% EtOH, then dried. Coverslips were incubated with 8.4mg/mL NaHCO3, 270 mg/mL mPEG-Succinimidyl Valerate, MW 2,000 (Laysan Bio), and 35 mg/mL Biotin-PEG-SVA, MW 5,000 (Laysan Bio) in a humid container for 3 hours. Slides were washed with water, dried, and stored at −20 °C or in a vacuum desiccator.

Taxol-stabilized microtubules with ∼10% biotin-tubulin and ∼10% Alexa488-labeled fluorescent-tubulin were prepared as previously described. Flow chambers were assembled with taxol-stabilized microtubules by incubating 1 mg/mL streptavidin in assay buffer (30 mM HEPES [pH 7.4], 450 mM potassium acetate, 2 mM magnesium acetate, 1 mM EGTA, 0.6 % methyl cellulose, 1 mM DTT) for 3 min, washing twice with assay buffer supplemented with taxol, incubating a fresh dilution of taxol-stabilized microtubules in assay buffer for 3 min, and washing twice with assay buffer supplemented with 1 mg/mL casein and 20 µM Taxol.

50 nM labeled KASH5 was incubated in the presence or absence of 100 nM KIF5B and 50nM MAP7 for 30 min on ice. They were then flowed into the flow chamber assembled with taxol-stabilized microtubules. The final imaging buffer contained the assay buffer supplemented with 20 µM Taxol, 1 mg/mL casein, 71.5 mM β-mercaptoethanol, 0.05 mg/mL glucose catalase, 1.2 mg/mL glucose oxidase, 0.4% glucose, and 2 mM AMP-PNP. Proteins were incubated with taxol-stabilized microtubules in the chamber for 10 min, followed by a wash with imaging buffer to remove unbound proteins. The KASH5-labeled sample was then imaged under single-molecule TIRF microscopy.

### Microtubule sedimentation assay

To further assess the potential interaction of KASH5 and KIF5B, taxol-stabilized microtubules were prepared similarly as mentioned above with unlabeled bovine brain tubulin. The 80 uL of polymerized microtubules were layered onto 100 uL of cushion buffer (80 mM PIPES [pH 6.8], 2 mM magnesium chloride, 1 mM EGTA, 1 mM DTT, 20 µM taxol, and 60 % [v/v] glycerol). The microtubule pellet fractions were collected by centrifugation (50,000 × rpm, 15 min, 37 °C) and resuspended in assay buffer (30 mM HEPES [pH 7.4], 2 mM magnesium acetate, 1 mM EGTA, 1 mM DTT, 20 µM taxol, 1 mg/mL casein, and 10% [v/v] glycerol). 200 nM KASH5 was incubated with 10 µM of the resuspended microtubules in the presence or absence of 40 nM KIF5B at room temperature for 20 min. The microtubules were then pelleted again by centrifugation (50,000 × g, 10 min, 26 °C). Supernatant and pellet fractions were collected and analyzed by SDS-PAGE.

### HeLa or Neuro 2 cell ectopic expression system

HeLa cells and plasmid constructs are described in (*54*). Cells were maintained in DMEM (Gibco) supplemented with 10% fetal bovine serum (FBS, Cytiva) and 1 % penicillin/streptomycin (Gibco) at 5.0 % CO_2_ and 37 °C. Cells were seeded onto cover slides in 24-well dishes at 40 % confluency. After overnight incubation, the DMEM media was removed and replaced with fresh DMEM containing 10% FBS without antibiotics. Cells were transfected with 0.5 µg of the SUN1-myc and FLAG-KASH5 (full length, FL, or lacking the trans membrane domain, ΔTM) constructs using Lipofectamine 2000 (Invitrogen) in Opti-MEM (Gibco) following the manufacturer’s instructions and incubated for 24 h.

The transfected HeLa or Neuro 2 cells were fixed/permeabilized in 4 % paraformaldehyde and 0.1 % Triton X-100 for 15 min at room temperature, then washed twice with 1 X PBS. The cells were then incubated in 3% bovine serum albumin (BSA) in 1X PBS for 1 hour at room temperature and washed with 1X PBS. Coverslides were incubated overnight at 4°C with primary antibodies: chicken anti-myc, mouse anti-FLAG, and rabbit anti-KIF5B, diluted in an antibody dilution buffer (0.1% Triton X-100 and 1% BSA in 1X PBS). Cells were then washed 3 times with 1X PBS before incubating with secondary antibodies diluted in the antibody dilution buffer for 1 h at room temperature in the dark. The following secondary antibodies were used: anti-chicken Cy5, anti-mouse TRITC, and anti-rabbit 488. After three washes in 1X PBS, the coverslips were mounted on glass slides with antifade mounting media containing DAPI (Vectashield). Slides were imaged using an LSM980 (Zeiss) equipped with an IR laser for multiphoton excitation confocal microscopy. All Z-stack images were acquired quantitatively with the same laser settings across experiments. Three biological replicates were performed, defined as three separate transfections. Line scans were generated in Z-projections of each image using the multiple plot profile tool from ImageJ, with the lines crossing the nucleus. ImageJ was used to generate inverted grayscale images, merge channels, and draw cell outlines, using contrast-adjusted images as needed. Some images were prepared and annotated using Adobe Photoshop.

### Immunofluorescence analysis in HeLa or Neuro 2cells

Analysis was performed using FIJI. Max projections of all channels were created. Cell boundaries were traced manually using the KIF5B channel as a marker for the cytoplasmic area. The cytoplasmic values were defined as the mean fluorescence intensity of the cell boundary minus a nuclear ROI expanded by 5 µm. The nuclear values of KIF5B and KASH5 were defined as the average fluorescence intensity in the corresponding channel, measured across five circular regions, each with a diameter of 0.35 µm, randomly distributed within the ROI that demarcated the nucleus. Cells were not considered if the KASH5 mean intensity fluorescent value was less than 200 for cells expressing KASH5-FL, or if the mean gray value of SUN1 or KASH5-ΔTM was less than 300. Data plotting and statistical analyses were performed in Prism9 (GraphPad). Only cells that were almost entirely in the field of view were analyzed.

### Co-immunoprecipitation experiments using HeLa or Neuro2 cells extract

KASH5 constructs used in this study: FL encoded full-length human KASH5, aa 1–562; ΔTM substitutes the TM domain aa 522–542 with a six aa gly-ser linker. For expression in HeLa cells or Neuro 2 cells, KASH5 constructs (FL and ΔTM) were cloned into a pcDNA3-derived vector with an N-terminal or C-terminal 3X FLAG tag. SUN1-FL cDNA was cloned into a pcDNA3-derived vector with a C-terminal 6X-myc tag. HeLa cells were transfected with 3 µg of SUN1 encoding plasmid alone (SUN1-myc) or together with KASH5-encoding plasmids (FLAG-KASH5-ΔTM or FLAG-KASH-FL) in a 6 cm dish. Transfections were performed with Lipofectamine 2000 (Invitrogen) following the manufacturer’s instructions. 24 h post-transfection, the media were removed, and the cells were harvested and washed with ice-cold 1X PBS. After centrifugation at 1000 X g for 2 min, cells were washed again with PBS and then transferred to microcentrifuge tubes for lysis. Lysis was performed by homogenization using 1 mL of ice-cold IP buffer (50 mM Tris-HCl, pH 7,5; 150 mM NaCl; 0,1 % IGEPAL; 0,5 % deoxycholate; 3 mM MgCl_2_; 3 mM EDTA; 10% glycerol; supplemented with protease and phosphatase inhibitors). After homogenization, the cell extract was kept on ice for 30 min. Lysates were then centrifuged at 50,000 X g for 20 min at 4 °C, and pellets were discarded. Input samples were collected. For immunoprecipitation, 2 μg of anti-KASH5, -KIF5B, or normal IgG antibodies were added to the cytoplasmic cell extracts, incubated overnight at 4 °C in rotation, followed by incubation with 50 μL of Dynabeads™ M-280 Sheep Anti-Mouse IgG (Invitrogen #11201D) for 1 h at 4 °C with rotation. Beads were washed four times with ice-cold IP buffer, and bound proteins were eluted by boiling for 5 min in SDS-PAGE sample buffer. Proteins were separated by 10% SDS-PAGE under reducing conditions and either analyzed by mass spectrometry or transferred to nitrocellulose membranes. The blots were probed with individual primary antibodies as indicated and then incubated with HRP-conjugated donkey anti-rabbit antibodies. In all blots, proteins were visualized by enhanced chemiluminescence. Two biological replicates were performed, defined as separate transfections and immunoprecipitations.

### Cell cycle analysis

After KASH5/SUN1 transfection (40% efficiency), HeLa cells were harvested by trypsinization, washed twice with cold PBS, and fixed with 4% PFA in PBS for 15 min. Next, cells were permeabilized with 0.1% Tween 20 in PBS for 10 min, then incubated with a mouse anti-FLAG antibody for 1 h at 37 °C, followed by a 45 min incubation with anti-mouse IgG-Alexa 488. Stained cells were washed with 1 X PBS and incubated with RNase A (100 μg/mL) for 30 min at 37 °C. DNA was stained with propidium iodide (PI; 50 μg/mL) for 30 min at room temperature in the dark. Samples were analyzed on a flow cytometer (BD FACSCanto II) using linear fluorescence detection for PI. At least 10,000 events were collected per sample, and doublets were excluded by gating on pulse width vs. area. DNA histograms were used to quantify the percentage of cells in G₀/G₁, S, and G₂/M. Cell cycle modeling was performed in FlowJo using the Dean–Jett–Fox algorithm. Percentages for each phase were compared between transfected (FITC+) and non-transfected (FITC-) cells.

### Spermatocyte squash preparation, immunostaining, and colocalization analysis

4-week-old mice testes were removed and detunicated, and the seminiferous tubules were processed for squashing. For squashing, we followed a technique previously described (*88*) with minor modifications. Briefly, seminiferous tubules were fixed in freshly prepared 2% formaldehyde in 1X PBS containing 0.1% Triton X-100. After 5 min, several seminiferous tubule fragments were placed on a slide and squashed using a coverslip; the coverslip was removed after freezing in liquid nitrogen. Samples were washed with 1X PBS and stored for up to 4 days before use.

We employed established experimental approaches to visualize chromosomes in spermatocyte squash preparations (*88, 89*). Incubations with primary antibodies were carried out for 12 h at 4 °C in 1X PBS with 2% BSA. To detect SYCP3, we generated a polyclonal chicken antibody raised against mouse SYCP3. KASH5 was detected using a polyclonal mouse antibody against mouse KASH5. KIF5B was detected using a rabbit antibody against mouse KIF5B. Tubulin was detected using a mouse antibody against mouse tubulin. After three washes in 1X PBS, slides were incubated for 1 h at room temperature with secondary antibodies. A combination of rhodamine-conjugated goat anti-mouse IgG with fluorescein isothiocyanate (FITC)-conjugated goat anti-rabbit IgG, and Cy5-conjugated goat anti-chicken IgG was used for simultaneous triple immunolabeling. Slides were mounted with Vectashield containing DAPI mounting solution (Vector Laboratories) and sealed with nail varnish. Quantification of co-localizing signals was performed by superimposing images from the corresponding fluorescent channels into a single-plane image. We use Axiovision SE 64 (Carl Zeiss, inc.) for imaging acquisition and processing. Statistical tests are described in the text or figures.

### Measuring RPMs in mouse spermatocytes

Acquisition and analysis of 3D time-lapse images in the mouse have been described previously (*11*). Before RPM measurements, seminiferous tubule explants (obtained from 6-8 weeks old mice) were incubated in DMEM media with 1 % DMSO-PBS (control) or the indicated kinesin inhibitors for 20 min at 32 °C before the assay.

Track speed mean is measured for each of 3-5 pericentromeric heterochromatic spots (revealed by Hoesch staining) per cell. The average speed of all quantified spots in one cell is reported. Because in the quantified cells spots do not fuse (merge) or fragment (split), then the average speed is given by the track length divided by the time between the first and last object in the track.

Track displacement length is the distance between the first and last spot’s position. The average track for all quantified spots (3–5) within a single cell is reported.

## Supporting information

All supplementary figures and tables

## Data Availability

The mass spectrometry data from this publication have been deposited to the Zenodo database [https://zenodo.org/record/7199989#.Y0tZGVLML1I] and assigned the identifier DOI: 10.5281/zenodo.7199989.

## Acknowledgments

This work was supported by NIH grants R21 HD103562, R01 GM125803, P30 NIH/NIGMS GM149376-01, and R01 HD110990 to P.R.J.; and ANPCyT-PICT 2019-1584 to B.C.G.; and ANPCyT-PICT 2020-00217 to D.Y; and R35 GM146739; R01 HD108809, and NSF 2142670 to D.M.E. The mass spectrometry experiments were supported by NIH grants R24 GM137786 and P20 GM103447 to KM. D.Y. and B.C.G. were supported by the Argentina National Council on Scientific and Technical Research (CONICET) and Fulbright-CONICET Fellowships. The authors greatly acknowledge the technical and imaging assistance of Dr. Cecilia Sampedro and Dr. Carlos Mas from the Centro de Micro y Nanoscopía de Córdoba – CEMINCO –CONICET–Universidad Nacional de Córdoba, Córdoba, Argentina.

## Supplementary Data

**Figure S1. Details of the assay to determine KASH5 and KIF5B domain interaction using direct yeast-two-hybrid analysis.** The schematic represents the predicted functional domains of KIF5B and KASH5 and truncation mutants created to map sites of KIF5B-KASH5 interaction.

**Figure S2. KASH5-SUN1 localizes with KIF5B at the nuclear envelope in HeLa or Neuro 2 cells.** Immunofluorescence of HeLa cells transiently transfected with SUN1-myc and FLAG-tagged KASH5-FL (A) or SUN1-myc and FLAG-tagged KASH5-ΔTM (B) constructs and analyzed using anti-KIF5B, anti-FLAG (KASH5), and anti-myc (SUN1) antibodies. Scale bar = 10 µm. (C) Transfected and control non-transfected HeLa cells were analyzed for DNA content by PI staining and flow cytometry. Representative DNA content histograms showing G₀/G₁, S, and G₂/M peaks in transfected (FITC+) or non-transfected (FITC-) samples are shown. (D) Immunofluorescence of Neuro 2 cells transiently transfected with SUN1-myc and FLAG-tagged KASH5-FL (top panel) or SUN1-myc and FLAG-tagged KASH5-ΔTM (lower panel) constructs and analyzed using anti-KIF5B and anti-FLAG (KASH5). Scale bar = 10 µm.

**Figure S3. KASH5/KIF5B interaction in HeLa and Neuro 2 cells.** (A) Western blots showing the results of immunoprecipitation using either a KIF5B antibody or an anti-FLAG (KASH5) antibody in HeLa cells, followed by immunoblotting with an anti-FLAG antibody (upper panel). The lower panel shows proteins coprecipitated with a KIF5B antibody or an anti-FLAG (KASH5) antibody, followed by immunoblotting with an anti-myc (SUN1) antibody. Inputs (1.5 % of total lysate) and molecular weight markers are also shown. (B) Western blots showing the results of immunoprecipitation using a KIF5B antibody in Neuro 2 cells, followed by immunoblotting with a KIF5B antibody and subsequently with an anti-FLAG (KASH5) antibody. Inputs (1.5 % of total lysate) and molecular weight markers are also shown.

**Figure S4. KIF5B inhibitor effect on the microtubule cell network and KASH5 loading at telomeres.** (A) Representative images of wild-type zygotene and pachytene spermatocytes immunostained with SYCP3 (marking chromosome axis), tubulin, and DAPI after incubation with 1 % DMSO-PBS and rose bengal lactone KIF5B inhibitor (5 µM). Experiments were repeated independently with spermatocytes obtained from three mice. (B) Assessment of KASH5 immunofluorescent foci formation in squashed seminiferous tubules treated or not with the KIF5B inhibitor. Quantitation of KASH5 foci per cell is also shown. Compared to control, 1% DMSO-PBS (30.2 ± 1.66 SUN1 foci per nuclei, n = 18 cells scored from spermatocytes obtained from 3 different mice), no significant reduction in the number of SUN1 foci apparently associated to the nuclear membrane were observed when seminiferous tubules were treated with 5 µM of rose bengal lactone (29.53 ± 1.6 foci per nuclei, two tailed t test P = 0.24, n = 19 cells).

**Table S1. Full list of microtubule-associated proteins obtained from mouse spermatocytes.** Processed, all: data containing the summatory for all signals of any individual protein found in any of the conditions studied. Processed, kin-dyn only: data containing the summatory for only kinesins and dyneins found in this screening.

**Table S2. Source and concentration of antibodies used in this study.**

