## Supplementary figures and images for "KIF5B drives meiotic chromosome dynamics via interaction with the KASH5-LINC complex"

### All supplementary figures and tables

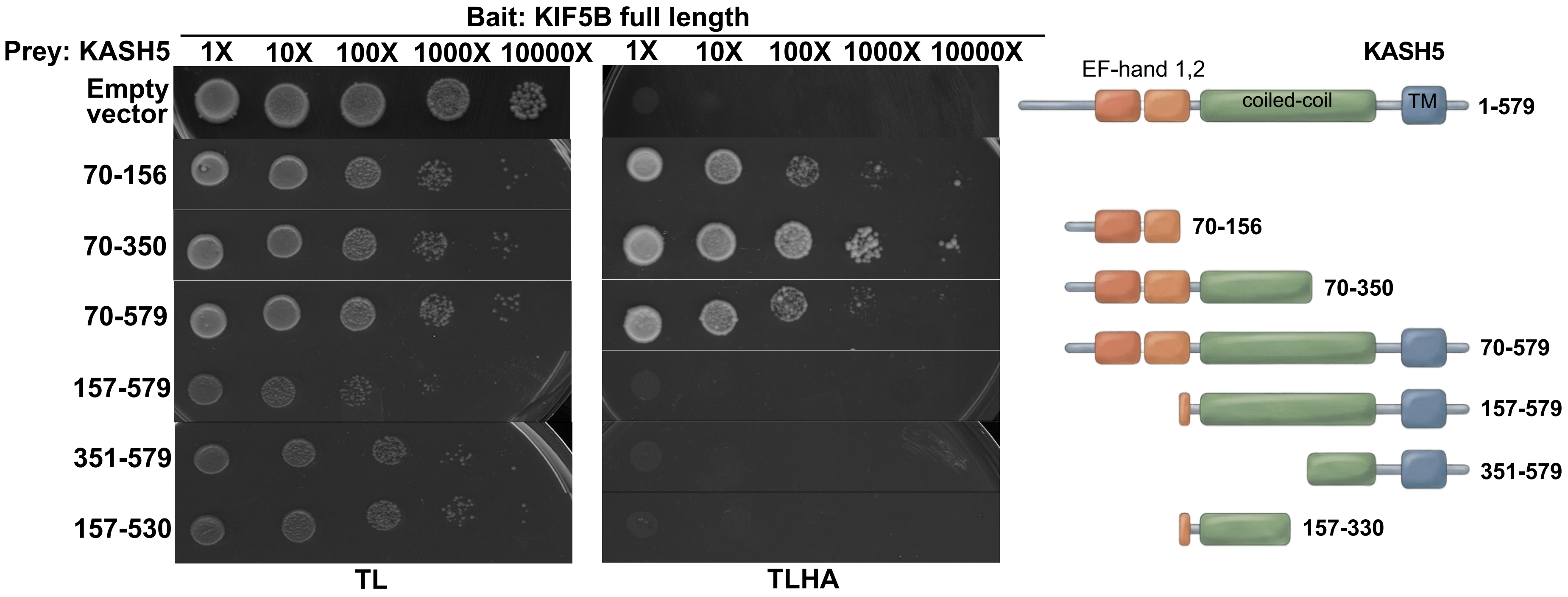

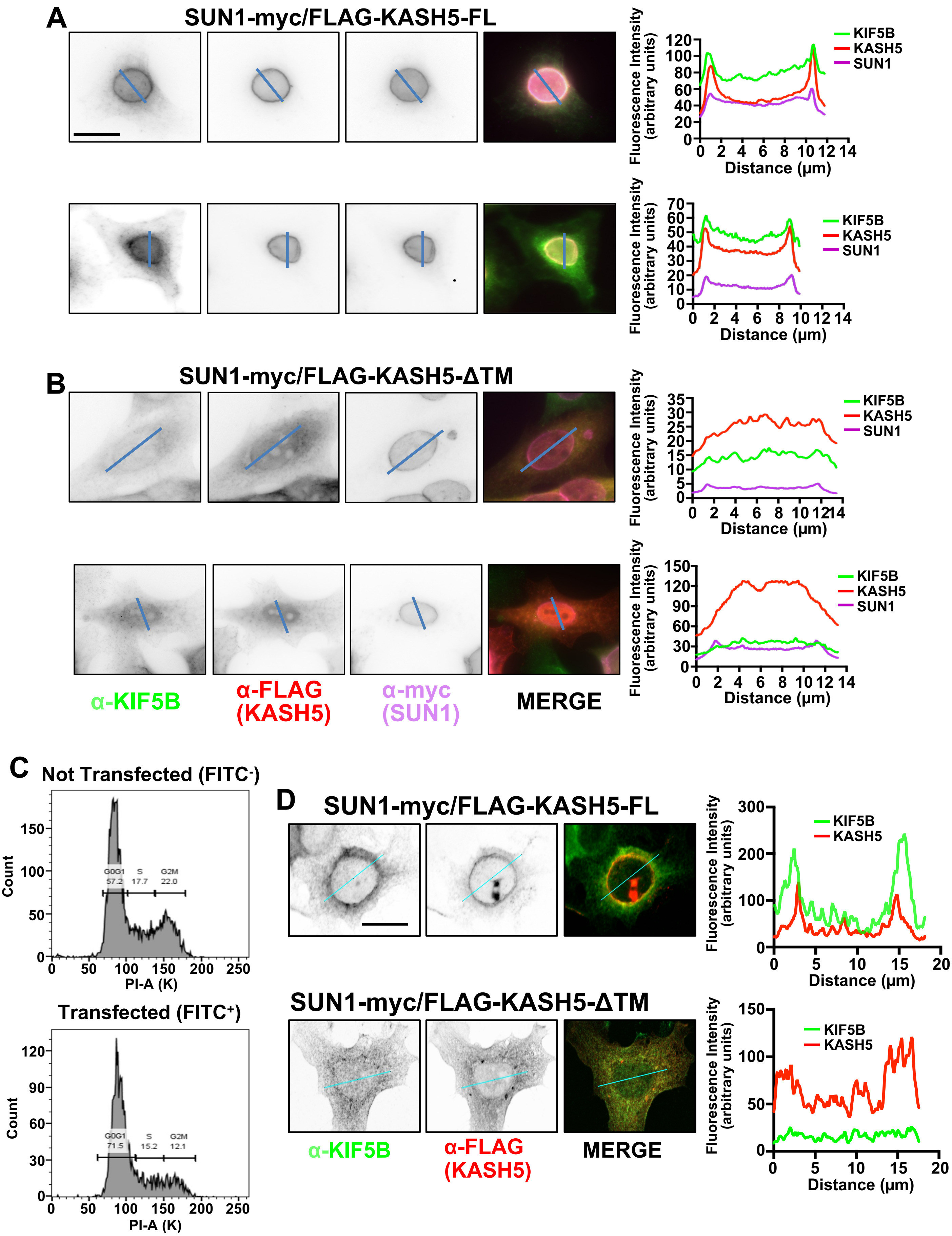

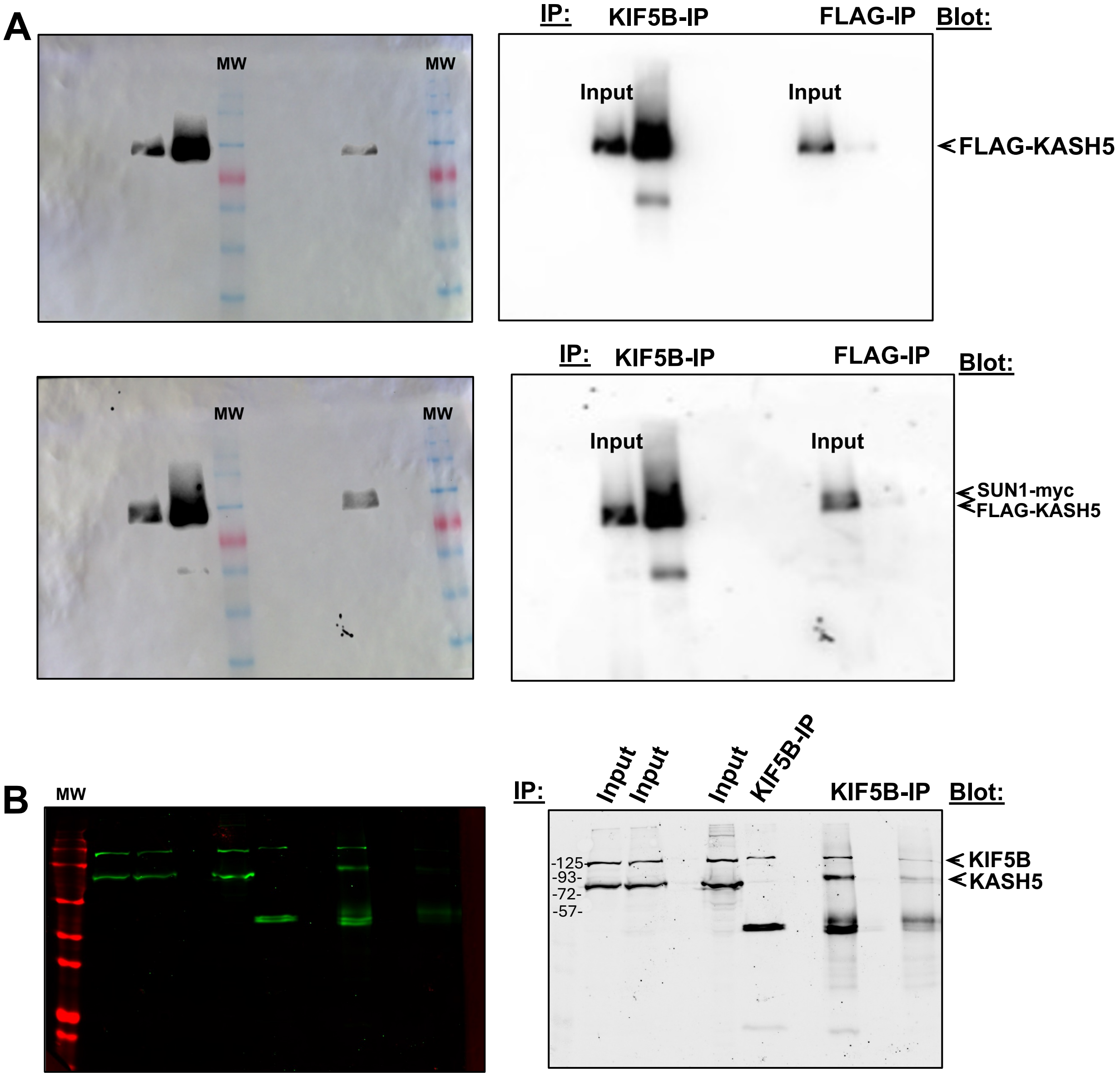

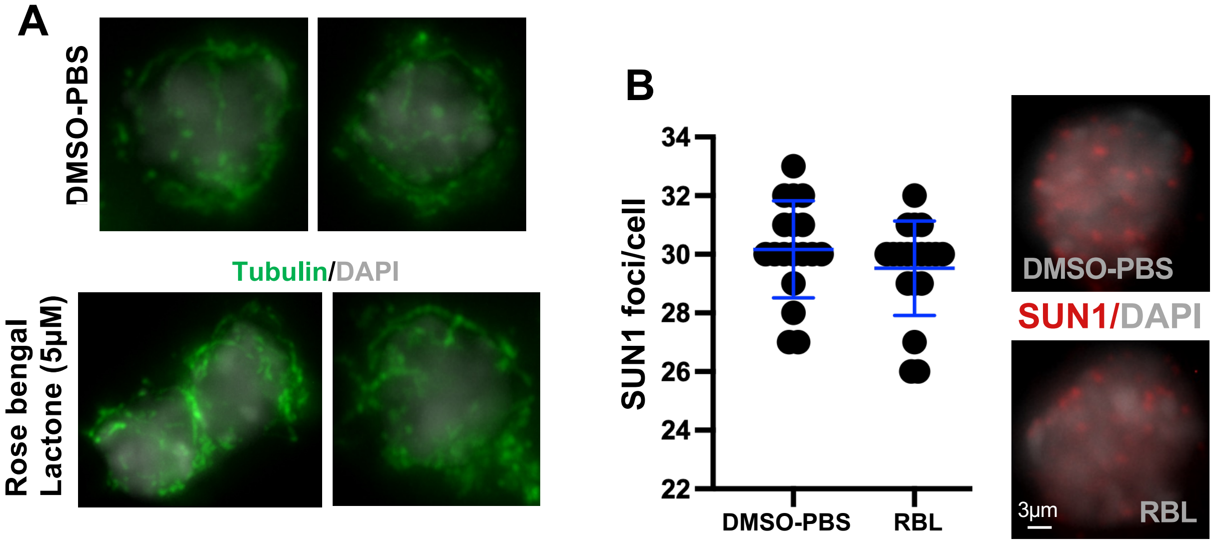
